# Handling ill-conditioned omics data with deep probabilistic models

**DOI:** 10.1101/2022.12.18.520909

**Authors:** María Martínez-García, Pablo M. Olmos

**Affiliations:** Signal Processing Group at Signal Theory and Communications Department, University Carlos III in Madrid (UC3M), Madrid, Spain; Gregorio Marañón Health Research Institute (IiSGM), Madrid, Spain

**Keywords:** Classification, Bayesian, dimensionality reduction, latent space model, missing data, VAE, omics, semi-supervised

## Abstract

The advent of high-throughput technologies has produced an increase in the dimensionality of omics datasets, which limits the application of machine learning methods due to the great unbalance between the number of observations and features. In this scenario, dimensionality reduction is essential to extract the relevant information within these datasets and project it in a low-dimensional space, and probabilistic latent space models are becoming popular given their capability to capture the underlying structure of the data as well as the uncertainty in the information. This article aims to provide a general classification and dimensionality reduction method based on deep latent space models that tackles two of the main problems that arise in omics datasets: the presence of missing data and the limited number of observations against the number of features. We propose a semi-supervised Bayesian latent space model that infers a low-dimensional embedding driven by the target label: the Deep Bayesian Logistic Regression (DBLR) model. During inference, the model also learns a global vector of weights that allows to make predictions given the low-dimensional embedding of the observations. Since this kind of datasets is prone to overfitting, we introduce an additional probabilistic regularization method based on the semi-supervised nature of the model. We compared the performance of the DBLR against several state-of-the-art methods for dimensionality reduction, both in synthetic and real datasets with different data types. The proposed model provides more informative low-dimensional representations, outperforms the baseline methods in classification and can naturally handle missing entries.

**Highlights:**

- *Inference of the latent space driven by the label value*. The DBLR infers different low-dimensional latent distributions depending on the label value, forcing clustering in the latent space in an informative manner, thus capturing the underlying structure of the data.
- *Classification*. During inference, the model additionally learns a global vector of weights that allows to make predictions given the low-dimensional representation of the data.
- *Handling missing data*. As the DBLR is a probabilistic generative model, it can naturally handle partially missing observations during the training process, including not annotated observations as censored samples. In this article we cover the Missing at Random (MAR) case.
- *Regularization method to handle small datasets*. In order to handle small high-dimensional datasets, which usually entail overfitting problems, we introduced an additional regularization mechanism following a drop-outlike strategy that relies in the generative semi-supervised nature of the model.
- *Handling different data types*. We have defined and implemented different observation likelihood models that can be used to describe different data types. In particular, we show how to use the DBLR with binary and real-valued features.

## 1. Introduction

As data acquisition methods evolve, scientific datasets grow rapidly in scale and complexity. This is particularly critical in omics, where the advent of high-throughput technologies has produced an increase in the dimensionality of the data, causing an important unbalance between the number of features and observations. This hampers the application of classification models on these datasets, as they are often limited by the *curse of dimensionality* [1]. Therefore, dimensionality reduction methods can be essential for further analysis in this scenario [30]. By means of different approaches, they learn a low-dimensional representation or embedding of the original data, which can be used for different applications as data visualization, data exploration or data classification.

Feature selection has been widely applied in bioinformatics to discard irrelevant features and reduce dimensionality in large datasets applying different information criteria. As of today, state-of-the-art in bioinformatics is given by the minimal-redundancy-maximal-relevance (mRMR) criterion [23] or the Hilbert–Schmidt Independence Criterion (HSIC) [8, 3]. Although these methods can result very useful for biomarker discovery, they do not infer a low dimensional embedding that captures the information within the data, but rather they select a subset of input variables.

Alternatively to feature extraction, linear dimensionality reduction methods such as Principal Component Analys (PCA) [10] or Canonical Correlation Analysis (CCA) [11] are widely used as one of the first steps in high-dimensional data analysis pipelines in order to overcome the limitations introduced by the large number of features, as the increased risk of overfitting or intractable computation [12, 26, 14, 15]. However, linear methods can fail to capture the complexity of high-dimensional datasets, which often imply non-linear dependencies within the data. For this reason, non-linear dimensionality reduction methods as t-Stochastic Neighborhood Embedding) (t-SNE) [18] or Uniform Manifold Approximation and Projection (UMAP) [19] have gained ground. In fact, UMAP has been rapidly adopted as dimensionality reduction method for preprocessing in genetic studies [4].

In this context, probabilistic latent space models have become increasingly popular for high-dimensional data analysis in recent years. Given their flexibility and capability to capture the underlying structures of the data, they can provide remarkable results in a variety of scenarios, and have proven to be particularly useful, for example, in cancer research, where the exploration of the latent space allowed the identification of cancer subtypes [25, 2, 9, 29, 28]. Additionally, the architecture of these kind of models can be easily adapted to different problems and types of input data, which have led to the development of a variety of methods for specific purposes such as netAE [5] or scVI [17], where the authors adapt the loss functions and architectures to properly analyze single cell RNASeq data, trying to maintain the local and global structure to capture cell relationships. However, these models are designed to be applied in a specific setup with a specific type of data, as expert knowledge is introduced in their definition.

In this work, we propose a general semi-supervised dimensionality reduction and classification method based on latent space models: the Deep Bayesian Logistic Regression (DBLR) model. Its main features, and thus our main contributions, are:

- *Inference of the latent space driven by the label value*. The DBLR infers different low-dimensional latent distributions depending on the label value, forcing clustering in the latent space in an informative manner, thus capturing the underlying structure of the data.
- *Classification*. During inference, the model additionally learns a global vector of weights that allows to make predictions given the low-dimensional representation of the data.
- *Handling missing data*. As the DBLR is a probabilistic generative model, it can naturally handle partially missing observations during the training process, including not annotated observations as censored samples. In this article we cover the MAR case.
- *Regularization method to handle small datasets*. In order to handle small high-dimensional datasets, which usually entail overfitting problems, we introduced an additional regularization mechanism following a drop-out-like strategy that relies in the generative semi-supervised nature of the model.
- *Handling different data types*. We have defined and implemented different observation likelihood models that can be used to describe different data types. In particular, we show how to use the DBLR with binary and real-valued features.

The structure of the paper is as follows. In Section 2 (Methods) we briefly introduce the Variational Autoencoder (VAE), which is the base of the proposed method. Then, we define the DBLR and explain the inference process, as well as the regularization of the model and how we handle MAR data. In Sections 3 (Results) we describe the experiments, introducing the baseline methods and the datasets, and show the obtained results. Finally, in Section 4 (Discussion) we summarise these results and draw the final conclusions of this work.

## 2. Methods

The DBLR is a semi-supervised Bayesian Logistic Regression model that follows a VAE [16] approach, learning a low-dimensional latent representation of high-dimensional data in order to classify it. In this section, we first briefly introduce the vanilla VAE, as it is the base of our proposed model. Then, we will describe in detail its definition of the DBLR, including the inference process, regularization and handling of MAR data.

### 2.1. VAE description

Autoencoders are Neural Networks (NNs) that learn a compact representation of the input data in a nonlinear unsupervised way, combining input features while minimizing the reconstruction error. In VAEs, this latent representation is learnt by means of Variational Inference (VI), assuming the data is coming from an underlying probability distribution. The goal is to fit the model parameters and to approximate the posterior distribution *p*(**z**|**x**), which describes the low-dimensional latent (encoded) representation **z** given the original (decoded) input data **x**. In the vanilla VAE, both the latent prior distribution *p*(**z**) and the likelihood *p*(**x**|**z**) are assumed Gaussian

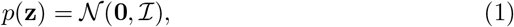

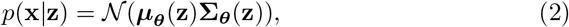

where the mean vector ***μ_θ_***(**z**) and diagonal covariance matrix **Σ*_θ_***(**z**) are the outputs of two NNs (decoder) with parameters ***θ***. Note the observations are i.i.d. under this model. Then, the marginal distribution *p*(**x**) is given by

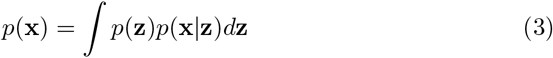

As this integral is intractable, it is not possible to do exact inference. Instead, the posterior distribution is approximated by means of VI assuming a Gaussian variational family

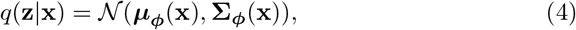

where the mean vector ***μ_ϕ_***(**z**) and diagonal covariance matrix **Σ*_φ_***(**z**) are the outputs of two NNs (encoder) with parameters *φ*.

The goal is to fit the parameters of the model to find the distribution that better approximates the posterior *p*(**z**|**x**) among the defined variational family by maximizing the Evidence Lower Bound (ELBO), which is given by

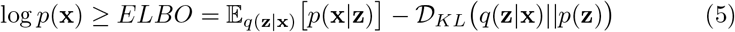

Using the negative ELBO as loss function for training, the likelihood of the input data (or reconstruction term) is maximized while keeping the approximated posterior close to the defined prior *p*(**z**), as the Kullback-Leibler (KL) divergence in the right acts as a regularization term.

### 2.2. DBLR description

The DBLR model generalizes the VAE in several ways. Assume a labelled 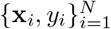 dataset where **x** is the D-dimensional feature vector and *y* is the target. We denote these variables as **X** = [**x**_1_, **x**_2_,…, **x***_N_*]*^T^* and **y** = [*y*_1_, *y*_2_,…, *y_N_*]*^T^*. We note that, although we assume binary labels, the model naturally handles categorical targets. As it can be seen in the graphical model depicted in Figure 1, one of the main differences of the DBLR is the assumption of dependency between the latent distribution and the target variable. This implies that the underlying structure of the data 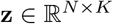, responsible for generating the properties of the observed samples **x**, also reflects the target variable. Thus, we force further clustering in the latent space in an informative manner, as the model captures the label information along the inference process ^1^. Additionally, following the idea of a Logistic Regression (LR) model, the DBLR learns a global vector of weights 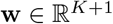 that allows to predict the label value given the lowdimensional representation of the input data. Table 1 summarizes the model variables.

**Figure 1:**
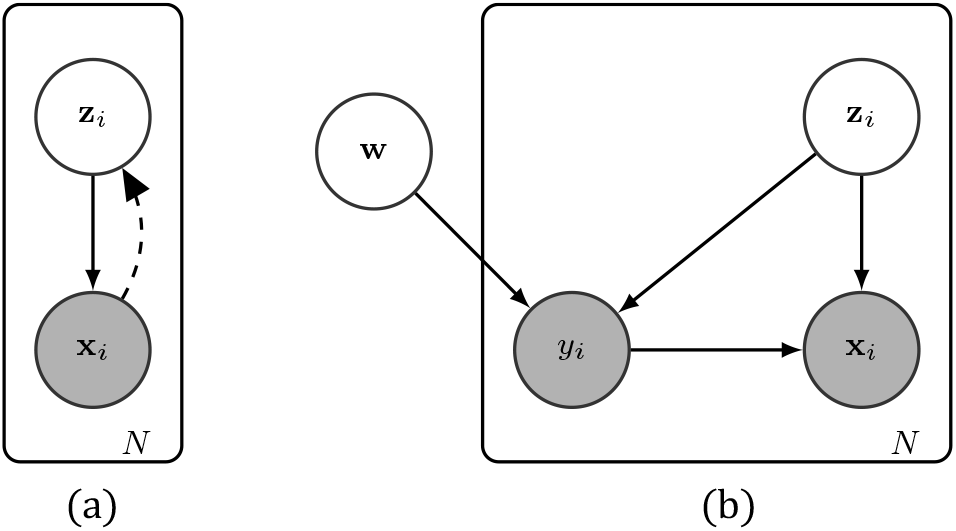
Graphical models. (a) Variational Autoencoder (VAE), (b) Deep Bayesian Logistic Regression model (DBLR)

**Table 1:** Description of the model variables.

|  |  |
| --- | --- |
| Input Data | $\mathbf{X} \in \mathbb{R}^{N \times D}$ or $\mathbf{X} \in \{0, 1\}^{N \times D}$ |
| Target Variable / Label | $\mathbf{y} \in \{0, 1\}^N$ |
| Low-dimensional Latent Variable | $\mathbf{Z} \in \mathbb{R}^{N \times K}, K \ll D$ |
| Global Weight Vector | $\mathbf{w} \in \mathbb{R}^{K+1}$ |

In the following sections we discuss the model in detail, including the inference process and the semi-supervised approach, used for handling not-annotated observations. We also introduce an additional regularization mechanism and explain how the model handles incomplete data.

#### 2.2.1. Prior distributions

We assume multivariate Gaussian distributions with zero mean and identity covariance matrix as priors for both global weights **w** and latent variable **Z**.

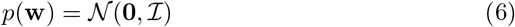

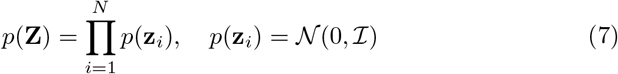

#### 2.2.2. Likelihoods

Depending on the nature of the input data, two different cases are defined:

- *Real input case* For the real input case, *p*(**X**|**Z, y**) is given by

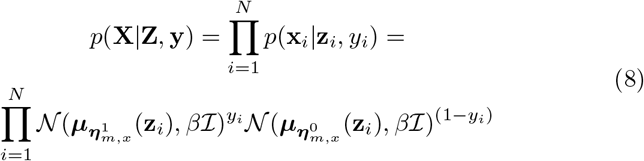

where the mean vectors ***μ*** are learnt by means of two NNs with input **z***_i_* and parameters 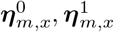. Superscripts indicate the value of the label, while subscripts indicate that the parameters correspond to the mean estimation NN. For simplicity of the model, we have selected a fixed diagonal covariance matrix with value *β*, which is an hyperparatemer of the model.
- *Binary input case* For the binary input case, *p*(**X**|**Z, y**) is given by

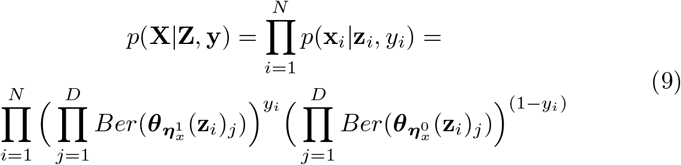

where ***θ**_**η**_x__*(**z***_i_*)*_j_* is the probability of feature *j* to be 1. These probabilities are computed by means of two NNs with input **z***_i_* and parameters 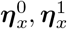. As in the previous case, superscripts indicate the value of the label. Observe that both in (8) and (9), the likelihood *p*(**x***_i_*|**z***_i_, y_i_*) critically depends on the target value, achieving a greater influence of *y* through the learning process. This way, we force the model to learn different parameters (and thus different distributions) for data with different target values, inducing clustering in the latent space. We also note that we could have a mixture between binary and real valued features. In such case, the term *p*(**x***_i_*|**z***_i_, y_i_*) would be the corresponding product of Gaussian and Bernoulli likelihoods.

Following the formulation of a LR model, the likelihood *p*(**y**|**Z, w**) is defined as

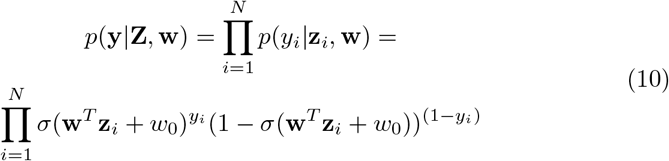

where *σ*(·) refers to the Sigmoid function. We introduce here a linear layer to keep the classification stage as simple as possible, forcing the model to infer highly separable latent representations to achieve a good classification performance. This way, the complexity relies in the generative model.

#### 2.2.3. Variational family and inference

The posterior distribution is approximated via VI, using the following variational family:

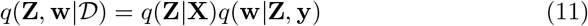

The first term, *q*(**Z**|**X**), which is given by

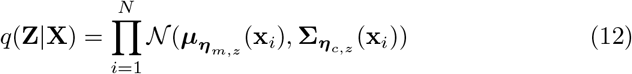

allows for inferring a latent representation of the observed data **X**. The mean vector and the diagonal of the covariance matrix are learnt by means of two NNs with parameters ***η**_m,z_* and ***η**_c,z_* respectively. Subscripts *m* and *c* stand for *mean* and *covariance*.

The second term, given by (13) refers to the posterior distribution of the weight vector **w**, which allows for classification. As it is a global variable, it is obtained as a product of the local contribution of each observed data point [22].

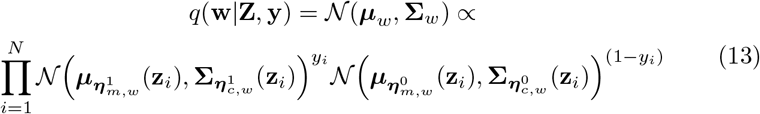

Mean vectors ***μ**_m,w_* and the diagonal of the covariance matrix **Σ***_m,w_* are learnt by means of four NNs with input **z**_i_ and parameters 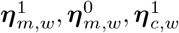 and 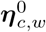. This way, we learn independent distributions or representations for the data with different label value.

Inference is done maximizing the Evidence Lower Bound (ELBO), which can be expressed as

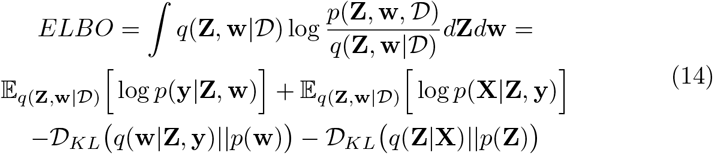

where the first two terms correspond to reconstruction of the label **y** and reconstruction of the input data **X**, respectively. The two remaining terms correspond to the KL Divergence between the variational family and the prior distributions. For further details, please refer to the appendix.

#### 2.2.4. Predictive distribution

To obtain predictions for new data, we first forward the new observation **x**\* through the encoder in (12) to obtain the parameters of the latent distribution. We combine (12) and (13) to derive the predictive distribution

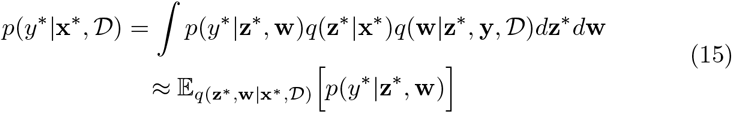

which we approximately evaluate using Monte Carlo sampling.

### 2.3. DBLR architecture

The architecture of the model can be divided in three main blocks: encoder, decoder and weight estimation block. Figure 2 shows a visual scheme of this architecture.

- *Encoder* The encoder consists of two fully connected NNs. One estimates the mean vector, and the other estimates the diagonal of the covariance matrix of *q*(**Z**|**X**), which is given by (12)
- *Decoder* The decoder consists of two fully connected NNs: one to learn the parameters for observations with *y_i_* = 1, and other to learn the parameter for observations with *y_i_* = 0. Depending on the nature of the data (real or binary), this NN estimates different parameters for *p*(**X**|**Z, y**). For the real input case, given by (8), it estimates a vector of means; for the binary input case, given by (9), it estimates a vector of probabilities.
- *Weight estimation block* This block consists of four fully connected NNs: two that learn the parameters of *q*(**w**|**Z, y**) (13) for observations with *y_i_* = 1, and other two for observations with *y_i_* = 0. Since we approximate the weights by means of Gaussian distributions, we learn a vector of weights and the diagonal of a covariance matrix for each label value.

**Figure 2:**
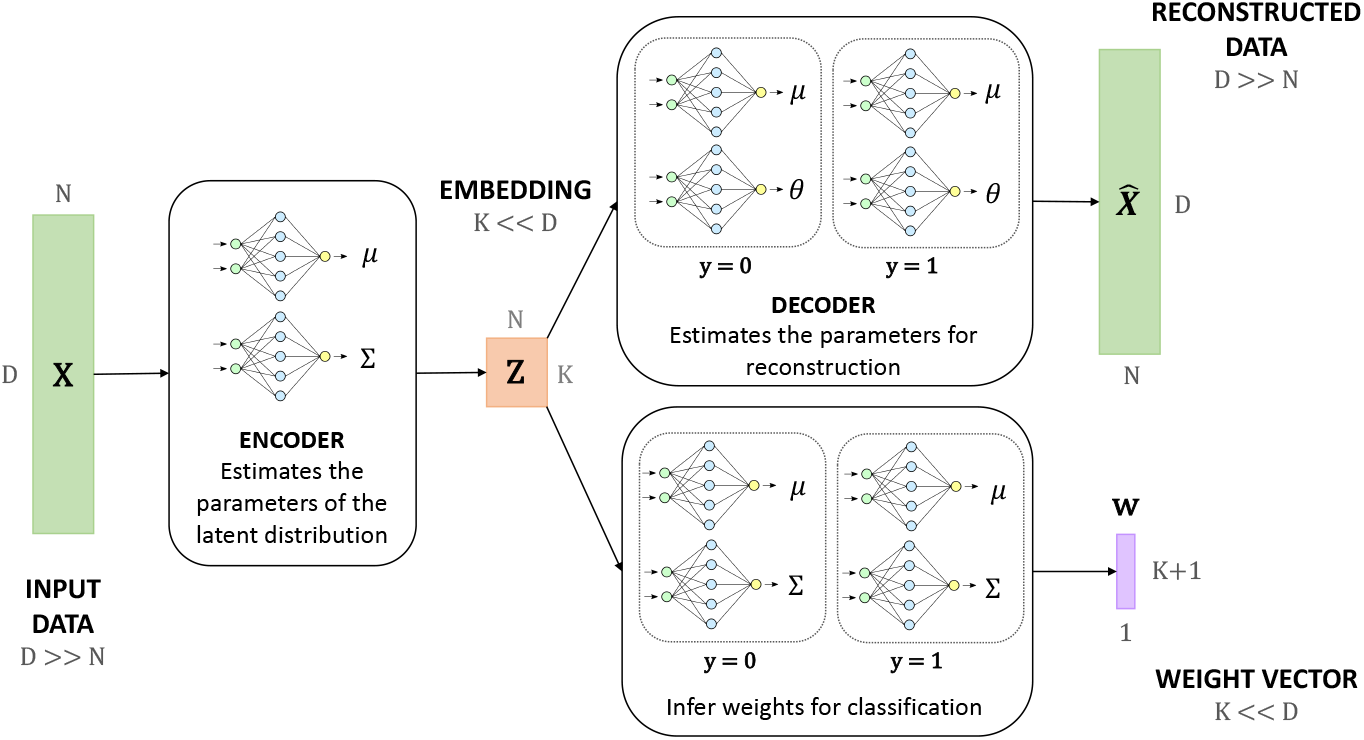
DBLR architecture.

### 2.4. Semi-supervised approach

As the presence of not-annotated observations or censored samples is a common problem in biological datasets, we define a semi-supervised approach to handle MAR data in the target variable *y*. When *y* is not known, we can evaluate the reconstruction term 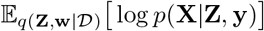 in the ELBO loss (14) by first sampling from *p*(**y**|**Z, w**), given by (10)^2^. In order to use gradient-based stochastic optimization, overcoming the inability to apply the reparameterization trick to discrete data, these samples are obtained by means of the Gumbel-Softmax trick [13]. Then, to evaluate the likelihood term 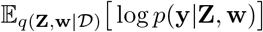 we only use observed y samples, i.e. we marginalize over the missing observations. This way, we only maximize the likelihood of the observed data.

### 2.5. Regularization

Although the model is already regularized by means of the KL Divergence terms present in the ELBO, which constrain the latent representation of the data, the model can still overfit since the reconstruction terms dominate over the KL terms. Thus, it is essential to introduce additional regularization mechanisms to achieve a compromise between the flexibility of the model and its generalization capability. In our case, we define a drop-out-like strategy that relies on the semi-supervised nature of the DBLR.

At each iteration of the gradient-descent optimization, we set a percentage *p* of the target **y** at missing. Then, the target value for these observations is sampled from *p*(**y**|**Z, w**) (10) by means of the Gumbel-Softmax trick, as the rest of the missing labels. Since this *censored* observations are selected at random, the model observes a different set of non-missing samples at each iteration of the learning process, preventing it from memorizing a given set of data.

### 2.6. Handling incomplete data

We have already described how the DBLR deals with not annotated samples, as it takes part in both inference and regularization processes. However, given its formulation, the DBLR can also handle partially missing observations in the feature vectors **x**.

If **x** is a vector with missing entries, to evaluate its projection over the **z** space using *q*(**z**_i_ |**x**_i_), we use the input-dropout trick described in [20], placing zeroes at missing entries in **X** and feeding the resulting vector 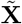. By placing zeroes, the output of the NNs that parameterize the distributions of the variational family are not affected by the missing entries. In Multilayer Perceptron (MLP) NN architectures, the outputs of the neurons are non-linear transformations of linear weighted sums of the inputs. Thus, the outputs and their derivatives do not depend on zero-valued entries. Then, the reconstruction terms in the ELBO loss (14) are only evaluated over observed attributes. Namely, we remove from 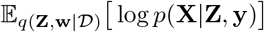 the factors corresponding to missing attributes.

## 3. Results

We compared our proposed model against different methods in synthetic and real high-dimensional datasets. Experimental details are explained along the following sections.

### 3.1. Baseline methods

In order to contrast the performance of our model against the state-of-the-art, we selected a set of different methods that are widely used for dimensionality reduction purposes in bioinformatics, making sure that we included models that use the target information. In particular, we compared our model against the following methods:

- *Supervised Uniform Manifold Approximation and Projection (S-UMAP)* [19]: it is a manifold learning algorithm constructed from a theoretical framework based in Riemannian geometry and algebraic topology. Although it was originally defined as an unsupervised dimensionality reduction method, it has been extended to include label information and thus obtain an embedding directly related with the target variable, as in our model.
- *Block Hilbert–Schmidt Independence Criterion (Block HSIC)*, [3]: it is a non-linear feature selection algorithm, which reduces the dimensionality of the input data based on the HSIC maximization between the high dimensional data and the resulting embedding. The *Block* version of the method reduces the memory complexity splitting the data in blocks. Note that this is a feature selection method, so it does not infer a low-dimensional representation of the data.
- *Principal Component Analysis (PCA)*, [10]: it solves the eigenvalue-eigenvector problem to generate a new reduced set of variables that attempts to explain the most variance of the data, while minimizing the residual variance.
- *Kernel Principal Component Analysis (KPCA)*, [27]: it is an extension of PCA that incorporates kernel methods. This enables the method to apply non-linear transformations to the data.
- *Canonical Correlation Analysis (CCA)*, [11]: it transforms the input variables to maximize the correlation between two datasets, which, in our case, correspond to the input features and the variables we want to predict.
- *Partial Least Squares (PLS)*, [7]: it generates a reduced set of components that attempts to explain as much as possible of the covariance between the input features and the target variable.

We used the Scikit-learn implementation for those methods available (PCA, KPCA, CCA and PLS). For UMAP and Block HSIC we used the implementation provided by the authors.^3^ ^4^

The goal of the proposed model is to reduce the dimensionality of the data inferring a latent distribution that captures its underlying structure while learning a vector of weights that enables classification of the observations. Thus, we compare the different methods in terms of the generated embeddings and in terms of classification performance.

For the classification experiment, we added a classification stage after the above mentioned methods by means of a LR including both L1 and L2 regularization. We decided to use a Logistic Regressor for this purpose to have a fair comparison between our model and the baselines, as the DBLR implements a Bayesian Logistic Regressor to describe the target variable and a non linear dimensionality reduction module. This way, we ensure the main differences in performance are mainly caused by the dimensionality reduction process and not by the characteristics of the classifier. However, it is worth mentioning that baseline methods were trained following a two-step approach where the latent representations were obtained prior to fitting the logistic regression model. In contrast, the DBLR optimizes the logistic regressor jointly with the latent representations.

For this experiment, we applied our model in two different manners. First, we trained the model over the train partition in a fully-supervised way, as the datasets do not present censored samples. We refer to these results as DBLR in the tables. Then, in order to exploit its semi-supervised nature, we trained the model over the full dataset setting test labels as missing. We refer to these results as DBLR SS in the tables (SS stands for Semi Supervised). However, it is important to remark that the semi-supervised capabilities of the model are also exploited in the DBLR version, since we are applying the previously defined regularization method based on internal censoring of targets at random.

Finally, we wanted to test the robustness of the model in the presence of partially-missing observations. For this purpose, we trained the model setting half of the training observations at missing for a given feature, with both samples and feature selected at random. Then, we compared the original observations against the imputation obtained by sampling from the model. Note this time we are introducing missing entries in **X**, while during the regularization process we are introducing missing entries in **y**. For simplicity, we use the cross-validated hyperparameters obtained in the previous experiment, since we are using the same data.

### 3.2. Datasets

- *Synthetic data* We generated synthetic data in order to compare the latent representations of different methods in a controlled setting. We start drawing samples from low-dimensional uniform mixture of Gaussian distributions with different mean vectors and covariance matrices, obtaining 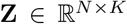. Samples drawn from the same Gaussian are assigned the same label. Then, we increase the dimensionality of the data applying a linear transformation by means of a random Gaussian matrix 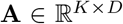. Assuming a binary dataset, we obtain **X** as

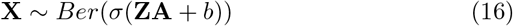

where 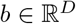 is a random bias vector.
- *Real data. The Cancer Genome Atlas (TCGA) dataset* Since the motivation behind the development of this method is to handle high dimensional datasets with limited number of observations, which are typical in omics, we analyzed four fat RNA-Seq datasets related to survival in different types of cancer. We used the tables preprocessed by [24], which contain RNA-Seq data extracted from the TCGA portal (https://tcga-data.nci.nih.gov/tcga/) and survival data extracted from the Genomic Data Commons (GDC) Portal (https://portal.gdc.cancer.gov/). They provide data for 32 different types of cancer, but we selected a subset (Breast Invasive Carcinoma (BRCA), Brain Lower Grade Glioma (LGG), Lung Adenocarcinoma (LUAD), Head and Neck Squamous Cell Carcinoma (HNSC)) based on the number of observations available. We additionally standardized the variables to have mean zero and unit standard deviation.

### 3.3. Hyperparameter tuning

The hyperparameters of the different dimensionality reduction methods have been tuned by means of 5-fold cross validation. We have validated the dimensionality of the embedding (the low dimensional representation of the data) for all the models except for CCA and Partial Least Squares (PLS), where the number of components is limited to *min*(*number of samples, number of features, number of classes*). As the target variables are binary, the number of components is limited to 2. More precisely, for each dataset we tried five different values for K, ranging from 2 to the total number of samples. In the case of our model, we have additionally validated the percentage *p* of samples we set at missing for regularization, trying four different values *p* = {0, 0.1, 0.25, 0.5}. Note that by setting *p* = 0 we are not introducing regularization. For the classification stage we have validated the regularization parameter, which we denote by *C* (for both L1 and L2 regularization), trying 10 different values that range from C = 10^-5^ to C = 10.

All the networks have been fixed to a basic fully-connected configuration which consists on one hidden layer of dimension either 25 or 100. This is the only network parameter we have validated. Note that cross-validation of other network parameters (number of layers, activation functions, dropout layers…) can potentially improve the experimental results.

### 3.4. Experimental results

In this section we present the results we have obtained applying the different baselines as well as our model to the datasets described above.

#### 3.4.1. Low-dimensional embedding on synthetic data

First, we analyze the low-dimensional representations generated by different methods when applied to a binary synthetic dataset. Since we generated high dimensional data drawing samples from a low-dimensional latent distribution, we can compare the embeddings learnt by some baseline models and the DBLR against the original latent space. We generated datasets of 500 samples and 10000 features following the generative process described in (16). Figure 3 shows the embeddings obtained applying the different methods, which were configured to generate latent representations of dimension *K* = 2 for proper visualization.

**Figure 3:**
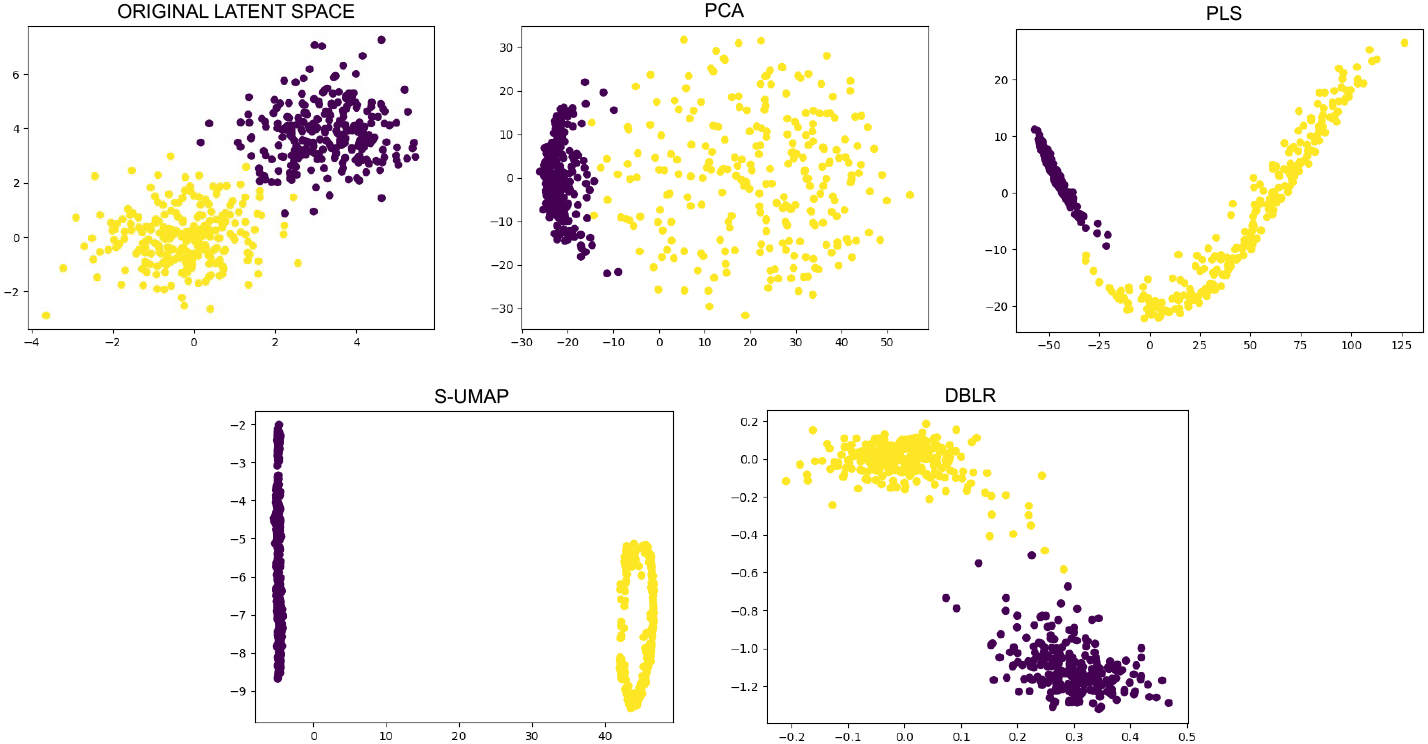
Low-dimensional embeddings generated by different methods against the original latent distribution for the binary input case (*N* = 500, *D* = 10000, *K* = 2).

Although all the methods generated highly separable low-dimensional representations that could be used for classification purposes, the obtained embeddings are quite different. While the DBLR preserves the natural structure of the data, the rest of the methods provide highly distorted and overfitted representations. The solution provided by the DBLR captures the original latent distribution of the data while enhancing the separation of the clusters, which reflects the label value.

#### 3.4.2. Classification on TCGA dataset

Now we analyze the classification performance of our model against the described baselines on real data using the Area Under the Receiver Operating Characteristics Curve (AUC). Since the number of observations is very limited in all the cases, we evaluate the performance of the different methods over 5 folds to avoid biases introduced by the partition of the data. For this reason, results are expressed in terms of mean and standard deviation. Notice that for each fold we treated train and test partitions independently.

Tables 3, 4, 6, 5 summarize the results obtained in classification for BRCA, LGG, LUAD and HNSC datasets, respectively. We show metrics obtained when applying the baseline methods as well as the DBLR in its basic and semisupervised form, referred as DBLR and DBLR SS in the tables. We include in the caption of every table the DBLR validated parameters. Namely, the latent dimension *K*, the networks hidden dimensionality (25 or 100), and regularization parameter p.

**Table 2:** Description of the four TCGA datasets.

| Dataset | # Observations | # Features |
| --- | --- | --- |
| BRCA (Breast Invasive Carcinoma) | 616 | 20221 |
| LGG (Brain Lower Grade Glioma) | 505 | 20223 |
| HNSC (Head and Neck Squamous Cell Carcinoma) | 473 | 20257 |
| LUAD (Lung Adenocarcinoma) | 434 | 20532 |

**Table 3:**
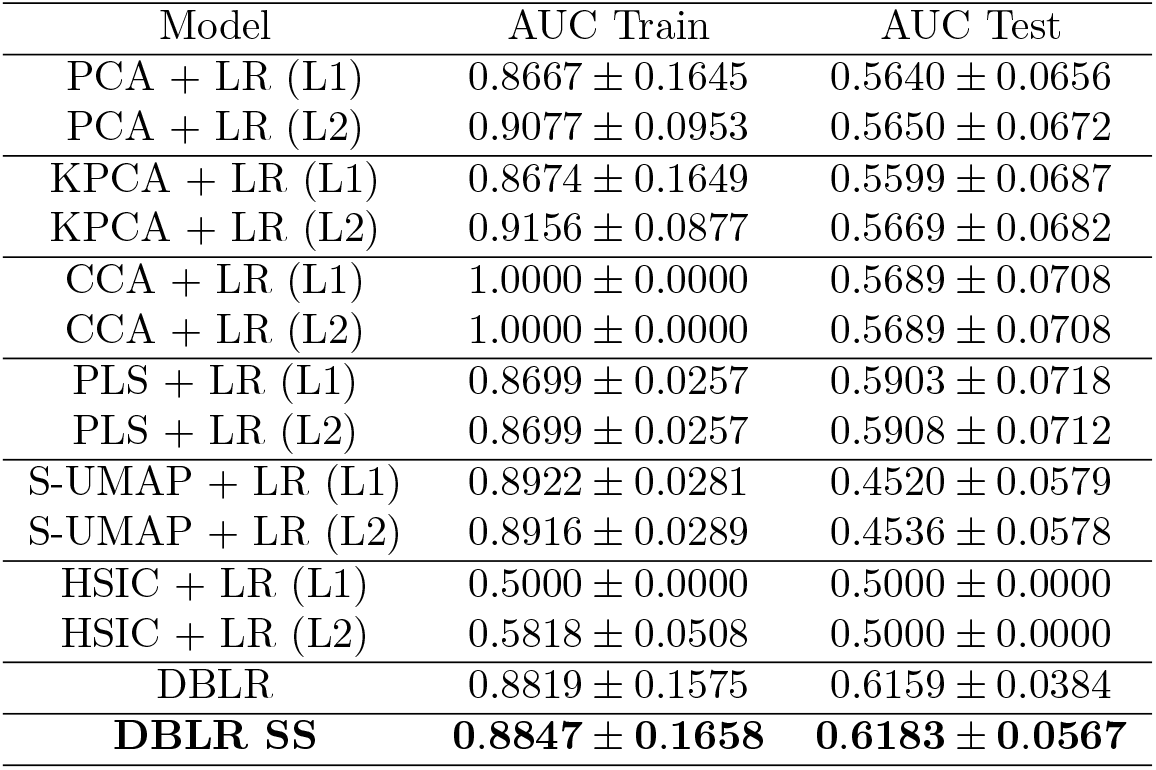
5-fold results for BRCA dataset. Mean values of the validated hyperparameters are *K* = 394.40, *p* = 0.06 and *hdim* = 25 for the DBLR, and *K* = 222.40, *p* = 0.27 and *hdim* = 100 for the semi-supervised DBLR.

**Table 4:**
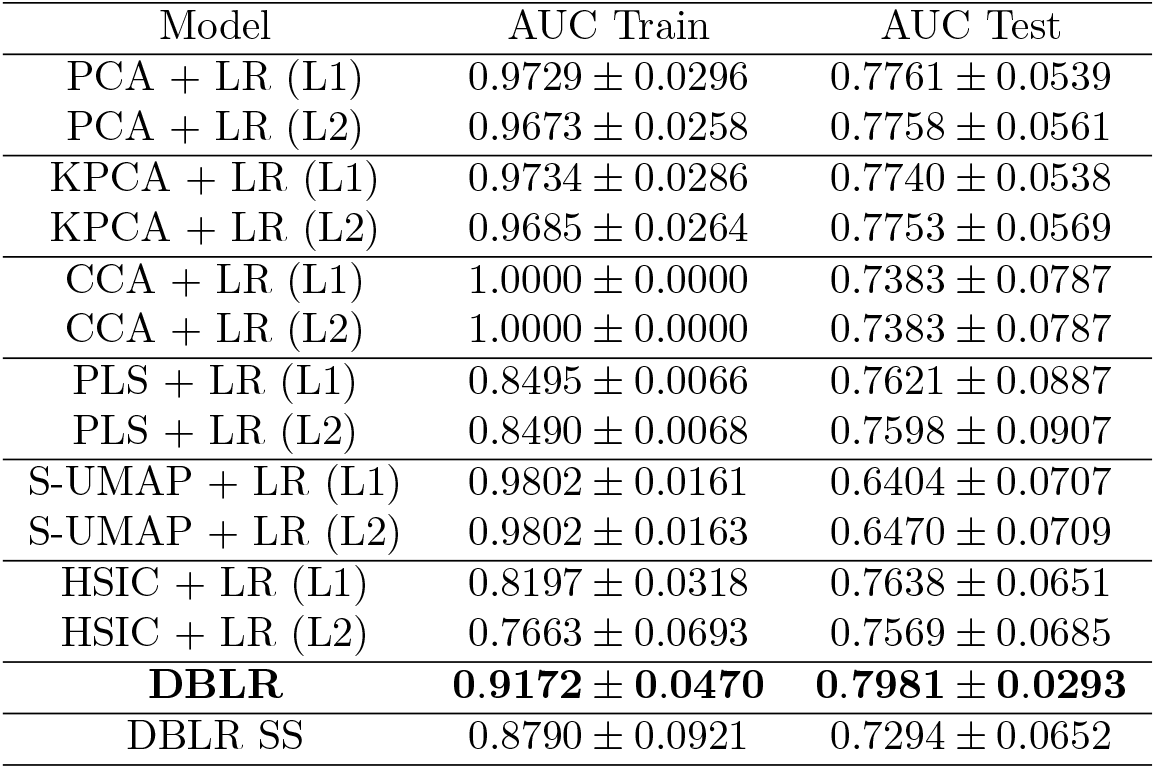
5-fold results for LGG dataset. Mean values of the validated hyperparameters are *K* = 303.20, *p* = 0.10 and *hdim* = 100 for the DBLR, and *K* = 263.20, *p* = 0.35 and *hdim* = 100 for the semi-supervised DBLR.

| Model | AUC Train | AUC Test |
| --- | --- | --- |
| PCA + LR (L1) | $0.9729 \pm 0.0296$ | $0.7761 \pm 0.0539$ |
| PCA + LR (L2) | $0.9673 \pm 0.0258$ | $0.7758 \pm 0.0561$ |
| KPCA + LR (L1) | $0.9734 \pm 0.0286$ | $0.7740 \pm 0.0538$ |
| KPCA + LR (L2) | $0.9685 \pm 0.0264$ | $0.7753 \pm 0.0569$ |
| CCA + LR (L1) | $1.0000 \pm 0.0000$ | $0.7383 \pm 0.0787$ |
| CCA + LR (L2) | $1.0000 \pm 0.0000$ | $0.7383 \pm 0.0787$ |
| PLS + LR (L1) | $0.8495 \pm 0.0066$ | $0.7621 \pm 0.0887$ |
| PLS + LR (L2) | $0.8490 \pm 0.0068$ | $0.7598 \pm 0.0907$ |
| S-UMAP + LR (L1) | $0.9802 \pm 0.0161$ | $0.6404 \pm 0.0707$ |
| S-UMAP + LR (L2) | $0.9802 \pm 0.0163$ | $0.6470 \pm 0.0709$ |
| HSIC + LR (L1) | $0.8197 \pm 0.0318$ | $0.7638 \pm 0.0651$ |
| HSIC + LR (L2) | $0.7663 \pm 0.0693$ | $0.7569 \pm 0.0685$ |
| **DBLR** | **$0.9172 \pm 0.0470$** | **$0.7981 \pm 0.0293$** |
| DBLR SS | $0.8790 \pm 0.0921$ | $0.7294 \pm 0.0652$ |

**Table 5:** 5-fold results for HNSC dataset. Mean values of the validated hyperparameters are *K* = 265.40, *p* = 0.17 and *hdim* = 25 for the DBLR, and *K* = 284.20, *p* = 0.04 and hdim = 100 for the semi-supervised DBLR.

| Model | AUC Train | AUC Test |
| --- | --- | --- |
| PCA + LR (L1) | $0.7941 \pm 0.0909$ | $0.6231 \pm 0.0510$ |
| PCA + LR (L2) | $0.7539 \pm 0.0197$ | $0.6340 \pm 0.0440$ |
| KPCA + LR (L1) | $0.7946 \pm 0.0921$ | $0.6238 \pm 0.0507$ |
| KPCA + LR (L2) | $0.7540 \pm 0.0198$ | $0.6340 \pm 0.0440$ |
| CCA + LR (L1) | $0.8347 \pm 0.0121$ | $0.6279 \pm 0.0348$ |
| CCA + LR (L2) | $0.8343 \pm 0.0125$ | $0.6265 \pm 0.0355$ |
| PLS + LR (L1) | $0.8347 \pm 0.0121$ | $0.6279 \pm 0.0348$ |
| PLS + LR (L2) | $0.8343 \pm 0.0125$ | $0.6265 \pm 0.0355$ |
| S-UMAP + LR (L1) | $1.0000 \pm 0.0000$ | $0.6038 \pm 0.0948$ |
| S-UMAP + LR (L2) | $1.0000 \pm 0.0000$ | $0.5792 \pm 0.1092$ |
| HSIC + LR (L1) | $0.7462 \pm 0.1438$ | $0.5526 \pm 0.0333$ |
| HSIC + LR (L2) | $0.6204 \pm 0.0375$ | $0.5415 \pm 0.0393$ |
| **DBLR** | **$0.7232 \pm 0.0884$** | **$0.6435 \pm 0.0365$** |
| DBLR SS | $0.9156 \pm 0.0520$ | $0.6014 \pm 0.0522$ |

**Table 6:** 5-fold results for LUAD dataset. Mean values of the validated hyperparameters are *K* = 19.20, *p* = 0.21 and *hdim* = 25 for the DBLR, and *K* = 225.80, *p* = 0.12 and *hdim* = 25 for the semi-supervised DBLR.

| Model | AUC Train | AUC Test |
| --- | --- | --- |
| PCA + LR (L1) | $0.6358 \pm 0.0257$ | $0.5853 \pm 0.0434$ |
| PCA + LR (L2) | $0.6842 \pm 0.0318$ | $0.6073 \pm 0.0543$ |
| KPCA + LR (L1) | $0.6358 \pm 0.0256$ | $0.5853 \pm 0.0433$ |
| KPCA + LR (L2) | $0.6843 \pm 0.0318$ | $0.6075 \pm 0.0541$ |
| CCA + LR (L1) | $1.0000 \pm 0.0000$ | $0.5521 \pm 0.0833$ |
| CCA + LR (L2) | $1.0000 \pm 0.0000$ | $0.5521 \pm 0.0833$ |
| PLS + LR (L1) | $0.8971 \pm 0.0118$ | $0.5805 \pm 0.0686$ |
| PLS + LR (L2) | $0.8971 \pm 0.0118$ | $0.5815 \pm 0.0691$ |
| S-UMAP + LR (L1) | $1.0000 \pm 0.0000$ | $0.5759 \pm 0.0521$ |
| S-UMAP + LR (L2) | $1.0000 \pm 0.0000$ | $0.6011 \pm 0.0458$ |
| HSIC + LR (L1) | $0.6832 \pm 0.1103$ | $0.6040 \pm 0.0690$ |
| **HSIC + LR (L2)** | **$0.6359 \pm 0.0097$** | **$0.6316 \pm 0.0539$** |
| DBLR | $0.6887 \pm 0.1307$ | $0.5994 \pm 0.0434$ |
| DBLR SS | $0.7867 \pm 0.1325$ | $0.6302 \pm 0.0308$ |

Given the classification results, we can say that the DBLR outperforms all baseline methods in BRCA (Table 3), LGG (Table 4) and HNSC (Table 5) datasets, and achieves the second best result in LUAD (Table 6) dataset. Note that in the latter, although the mean test AUC obtained with the DBLR is slightly worse than the one provided by the HSIC, its standard deviation is lower, meaning that the results are more stable.

In the case of LGG and HNSC, the basic version of the model outperforms the semi-supervised approach, while for BRCA and LUAD, the semi-supervised version achieves the best performance. These results may suggest that not all the datasets benefit from the semi-supervised approach, and one possible explanation is the additional introduction of noise through not annotated samples (recall that we are handling test samples as the target was missing). However, some datasets may benefit from this approach, since the introduction of these not annotated observations can additionally regularize the training process and thus, improve generalization.

If we compare the performance of the different baselines, we can see that S-UMAP achieves the worst or one of the worst classification metrics for all the datasets in test. Linking these results with the ones obtained for synthetic data, we can hypothesise that this dimensionality reduction method tends to provide overfitted solutions that may not be suitable for further analysis, although it can be a powerful tool for data visualization.

In the case of the HSIC, the results vary importantly across the different datasets. As this is a feature selection method, results depend on the presence or absence of variables that are highly correlated with the target. The rest of the methods can combine input features to extract new variables that capture the structure of the data, but in this case we can only select the ones that provide more information to the problem, but it may not be enough in all the scenarios.

For Kernel Principal Component Analysis (KPCA) and PCA, we obtained very similar results in all datasets. We trained the KPCA using a nonlinear kernel, particularly the Radial Basis Function (RBF) kernel. However, the parameter *γ*, which indicates the inverse of the standard deviation of the RBF kernel, was set to zero during hyperparameter tuning in all the cases, which is equivalent to use a linear kernel. This implies that both methods are applying a linear dimensionality reduction and thus, they achieve similar results.

It is important to confirm that our proposed method can achieve state-of-the-art classification performance while inferring a low-dimensional latent distribution of the data, capturing its underlying structure. In addition, given the probabilistic generative nature of the DBLR model, it can handle partially missing observations in the feature vectors and the target variable.

#### 3.4.3. Imputation of MAR data on The Cancer Genome Atlas (TCGA) dataset

Finally, we show that the DBLR can be trained with partially-missing observations, which can be imputed sampling from the model. Note that neither of the baseline methods can naturally deal with partially observed values or allow their probabilistic imputation. Figure 4 shows the imputation results obtained for a random feature in LGG dataset. Half the observations of that feature were set to missing in the train partition. Once the model was trained, we projected the samples over the latent space using *q*(**z***_i_*|**x***_i_*) in (12) and then reconstructed the low-dimensional representation using *p*(**x***_i_*|**z***_i_*, **y***_i_*) in (8), obtaining an imputation of the missing values. We show the results for both the observed and missing samples in train, as well as the results for the test partition, where we treated all the samples as missing entries. This way, we show that the model can impute missing values of new observations not used during training. Additionally, we compare our results with the ones obtained using a K-Nearest Neighbors (KNN) imputer ^5^ and sampling from a normal distribution with the empirical mean and variance of the observed training data.

**Figure 4:**
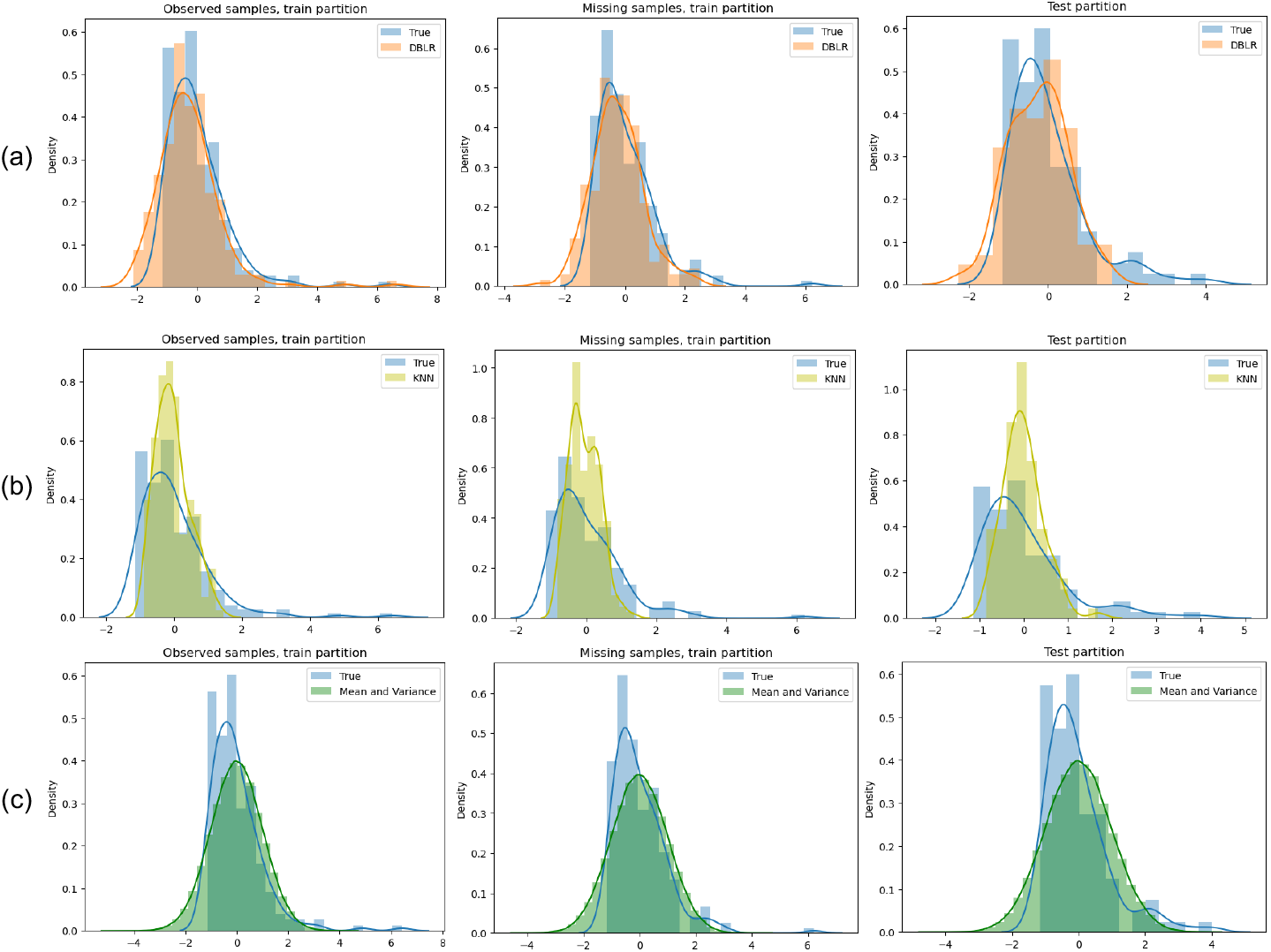
Empirical distribution of the data (represented in blue) against the inferred distributions obtained applying different methods. (a) Results using the DBLR. (b) Results using a KNN imputer with 5 neighbors. (c) Results using the empirical mean and variance of the observed training data.

The results show that the model is robust against missing entries, as it can still provide a good approximation of the distribution of the data. In fact, the distribution inferred by the DBLR is the one that better approximates the true data, capturing both the mode and the tails of the true density. The results show that the KNN imputer does not capture the tails of the distribution and the Normal distribution with the empirical mean and variance fails to capture the mode of the data.

## 4. Discussion

In this work, we presented the DBLR, a semi-supervised bayesian latent space model that follows a VAE approach to infer a low-dimensional representation of high-dimensional input data. The inference process is driven by the label value, which allows to capture the underlying structure of the data and thus obtain a separable latent representation. Additionally, the model learns a global vector of weights to predict the label value given the low-dimensional embedding of the input data.

We showed in both synthetic and real databases that the model can infer informative embeddings, capturing the nature of the latent distribution of the data while enhancing separation with respect to the label value. This separation in the latent space allows the DBLR to obtain better classification results than other state-of-the-art methods given the low-dimensional embedding, as showed in the experiments.

In this context, it is important to remark that the proposed method is a probabilistic generative model. This implies that it infers probability densities we can draw samples from. Thus, it can handle missing entries in both the target variable and the input data, which is particularly relevant in clinical and life-science scenarios, where it is common to find censored samples and partially missing observations. We showed that the DBLR can be trained in the presence of missing values and still infer the distribution of the data (for both observed and missing entries).

The proposed model not only can handle MAR data, but it benefits from this capacity to additionally regularize the training process by means of a drop-out-like strategy: a percentage of the observed targets is set as missing randomly and their values are imputed sampling from the model. This way, the model learns from a different set of observed samples at each training epoch.

## Appendix A. DBLR description

### Appendix A.1. Variational Family

As it is described in the main document, the posterior distribution 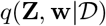 is approximated via VI, using the following variational family:

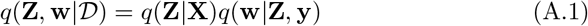

The second term, which refers to the posterior distribution of the weight vector **w**, is given by

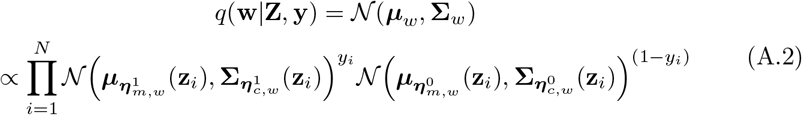

where the mean vector is given by

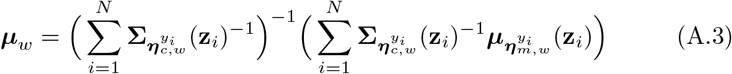

and the covariance is computed as

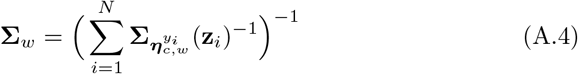

Recall that, as it is a global variable, it is obtained as a product of the local contribution of each observed data point. The mean vectors ***μ**_m,w_* and the diagonal of the covariance matrix **Σ***_m,w_* are learnt by means of four neural networks with input **z***_i_* and parameters 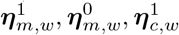 and 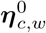.

### Appendix A.2. ELBO

Inference is done maximizing the Evidence Lower Bound (ELBO), which can be expressed as

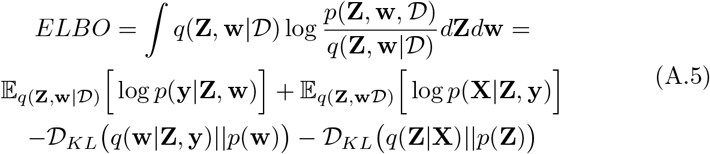

The first term (A.6) corresponds to the reconstruction of the label **y**, where *σ*(·) refers to the Sigmoid function.

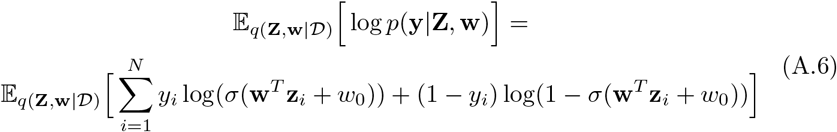

The second term corresponds to the reconstruction of the input data. Maximizing this term, we are inferring the low dimensional representation of the data that allows for the best possible reconstruction of **X**. Depending on the nature of the data (*real* (A.7) *orbinary* (A.8)), this term can be expressed as

- *Real input case*

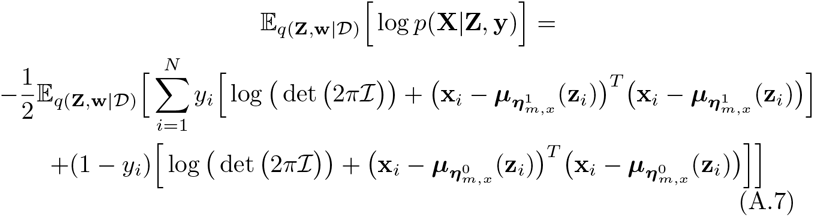
- *Binary input case*

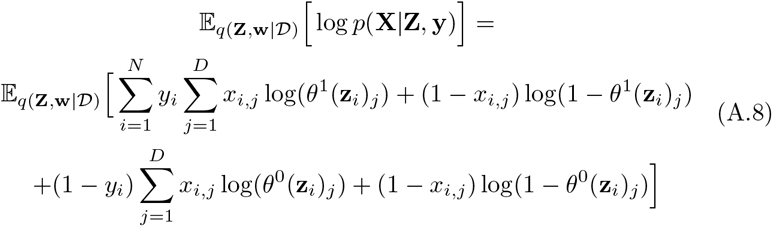

The two remaining terms allow for regularization. These correspond to the KL Divergence between the variational family and the prior distributions. Since the priors are Gaussian, the KL Divergences can be expressed in closed form as

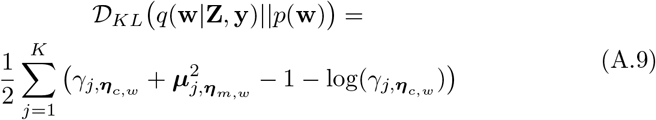

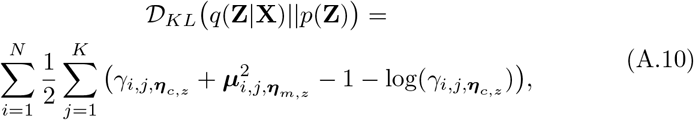

where *γ* refers to the diagonal of the covariance matrix **Σ**.

Although the reconstruction term given by (A.7) and (A.8) may numerically dominate over the likelihood term (A.6) in the case of high-dimensional inputs, the model can still learn to make accurate predictions, as the target variable is indirectly used during inference and regularization. This means that prediction of label values is embedded in the computation of the rest of the ELBO terms, so the model can learn to properly predict despite the possible unbalance between the different likelihood terms.

## Footnotes

1 This is common to other supervised VAE-like models such as [31, 32, 6, 21]

2 This is equivalent to assume an augmented variational family *q*(**Z**|**X**)*q*(**w**|**X***_o_*, **y***_o_*)*p*(**y***_m_*|**Z, w**), where *o* is the index set of observed targets and *m* the set of not observed ones.

3 UMAP package documentation at https://umap-learn.readthedocs.io/en/latest/basic_usage.html

4 Block HSIC implementation at https://github.com/riken-aip/pyHSICLasso

5 We used Scikit-learn implementation.

